# Parental care shapes evolution of aposematism and provides lifelong protection against predators

**DOI:** 10.1101/644864

**Authors:** C. Lindstedt, G. Boncoraglio, S.C. Cotter, J.D.J. Gilbert, R.M Kilner

## Abstract

Social interactions within species can modulate the response to selection and determine the extent of evolutionary change. Yet relatively little work has determined whether the social environment can influence the evolution of traits that are selected by interactions with other species - a major source of natural selection. Here we show that the amount of parental care received as an offspring can influence the expression, and potential evolution, of warning displays deployed against predators in adulthood. In theory, warning displays by prey are selected by predators for uniformity and to reliably advertise the extent to which individuals are chemically defended. However, the correlated evolution of intensity of the visual display and strength of the chemical defense is only possible if there is a genetic correlation between them. Adult burying beetles *Nicrophorus vespilloides* bear bright orange elytral markings which advertise their chemical defenses. We experimentally manipulated the level of maternal care that individuals received when they were larvae and then measured the strength of the correlation between the component parts of the warning display when they reached adulthood. We found that under limited care individuals were smaller and produced less conspicuous warning displays. The underlying family (genetic) correlation between the visual display and the chemical defense was weaker in individuals that received little care as larvae. We conclude that parenting by burying beetles modulates the evolvability of aposematic defense, making correlated evolutionary change in signal intensity and chemical defense less likely when they restrict care to their young.

**Significance Statement:** Parental care can improve early offspring survival against predators. However, we have little knowledge of how its effects shape the evolution of predator-prey interactions later in the offspring’s life. We tested this with carrion beetles who provide care for offspring and who carry warning coloration to advertise to predators that they are chemically defended. We show that more parental care resulted in larger, more brightly coloured and chemically defended adult beetles. Furthermore, when parents had provided little care for their young we found weaker genetic correlations between warning signal salience and chemical defense. Over time, this could result in untrustworthy warning signals, which could render them ineffective against predators.

## INTRODUCTION

Social interactions among conspecifics constitute an individual’s social environment. The social environment imposes selection directly on the traits that mediate social interactions within species such as the maintenance of honest signaling in quorum sensing bacteria (1). The social environment can also influence evolution more indirectly by accelerating or preventing a response to directional selection (2, 3) and by changing the adaptive value of the traits it induces (4, 5). In previous work, the influence of the social environment on trait evolution has been analysed mainly for traits that are selected by social interactions within species, such as body size (which influences competitive ability) or signals between conspecifics (3, 6–8). However, interactions with other species, such as predator-prey or host-parasite, are also an important source of selection. Relatively little work has determined how the social environment *within* species might influence the evolution of traits under selection from a different species (9–11).

Here we address this neglected issue by analysing how the level of parental care received during development might influence the evolution of antipredator and signal traits involved in predator-prey interactions later on in life, during adulthood. The level of care supplied by parents is well known to be influenced by diverse intraspecific interactions including sexual conflict between parents, conflict between parents and offspring and competition and cooperation between offspring over resource division (2, 6, 12–14). Since interactions within the family play out during development, they can have a major non-genetic role in determining an individual’s phenotype (3, 4, 6, 14). Importantly, these effects can persist into adulthood and can shape an individual’s fitness throughout its lifespan (e.g. (4)). However, it is much less well understood whether the effects of parent-offspring interactions have similarly enduring effects on traits involved in ecological interactions (15, 16). Specifically, it is unclear whether the social environment created by conspecifics can influence the extent of phenotypic variation that is exposed to selection from other species, due to predation (17). Yet this is key to understanding the evolutionary dynamics of the traits that mediate such interspecific interactions, particularly if it means that the extent of phenotypic variation then differs from the underlying genetic variation.

We tested this with the aposematic burying beetle (Fig S1), *Nicrophorus vespilloides*, an insect that exhibits biparental care (18). Parents breed on the carcass of a small vertebrate, like a mouse or a songbird, which they convert into an edible nest for their young (19). They bury it below ground, shave off the fur or feathers, roll the flesh into a ball and cover it in antimicrobial fluids (20). After the larvae hatch, the parents stay with their young to defend the carrion and their offspring from attackers, and they also feed their larvae via oral trophallaxis (19). There is considerable natural variation in the duration of post-hatching care among families (21–23). Some broods receive no post-hatching care at all but some larvae can nevertheless survive (6).

Adult burying beetles bear orange and black coloration (Fig. S1), which is used to communicate their chemical defences to predators (18). When handled, they produce a droplet of foul smelling fluid from their abdomen (18). In general, the toxic defences of an aposematic individual are considered to contribute to a ‘public good’ by educating predators to avoid prey bearing a similar signal (24). Through this public goods system of education, predators are predicted to select for uniformity and reliability in warning displays (25–27). Furthermore, greater levels of toxicity (28, 29) and pronounced signal patterns (30, 31) are expected to evolve under directional correlated selection by predators, as all these traits enhance the avoidance learning rate of predators by making warning displays memorable and easy to recognize.

We tested whether experimentally manipulating the supply of parental care for the larvae 1) caused a change in the aposematic display and if it did, 2) whether care *independently* affected the two elements of the aposematic display (the visual warning component and the chemical defence) when larvae matured into adults. We predicted that the supply of parental care potentially affects phenotypic variation in warning display in two different ways. First, it may fund the cost of producing toxic chemical defences and the pigmentation used in the warning signal. Individuals under full care should have better access to resources during the larval stage (32, 33). As a result, their warning signals should be larger and more salient, and chemical defences should also be more repellent.

The second way in which the supply of care could influence the evolution of warning signals is by causing phenotypic variation in the extent to which the signal accurately indicates the potency of an insect’s chemical defences (34). A scenario like this might arise if the costs of producing chemical defences and coloration are not equal, for example, or if they draw unequally on resources supplied by parents. This is an extension of a previous suggestion, that the extent of covariation between the salience of a warning signal and its associated toxicity is determined environmentally (34–36). However, for this type of trade-off to occur, there needs to be either a physiological linkage (e.g. constrained amount of antioxidants are needed for both bright pigmentation and protection of tissues against autotoxicity (35)) or genetic correlation between signal salience and chemical defence. If this correlation is weak then a reliable warning signal is less likely to evolve in response to selection from predators. Conversely, if this correlation is strong, we can expect correlated responses to selection from predators in both chemical defence and signal conspicuousness. To test this second prediction, we measured the broad sense heritabilities of each trait and their family-level (genetic) correlations across treatment groups with varying amounts of parental care to test whether the social environment can alter the co-evolvability of signal salience and levels of chemical defence.

## RESULTS

### Body size

We manipulated the level of post-hatching parental care, by withdrawing parents at different intervals after larval hatching in four different experimental treatments (0h, 8h, 24h, 48h). In general, we found that parental care affected the quality of the offspring. The greater the duration of care, the greater the adult body mass attained by offspring of both sexes (Care duration: F_1,94_ = 17.19, P < 0.001; Figure 1a), and the larger the elytra (Care duration: F_1,48_ = 8.08, P = 0.006; Figure S2b). Elytra size did not differ between the sexes, but males were significantly heavier than females (Sex: F_1,737_ = 14.5, P < 0.001, Fig. 1a). The interaction between care duration and sex was not significant for either trait (F<2.03, P>0.15).

**Figure 1.**
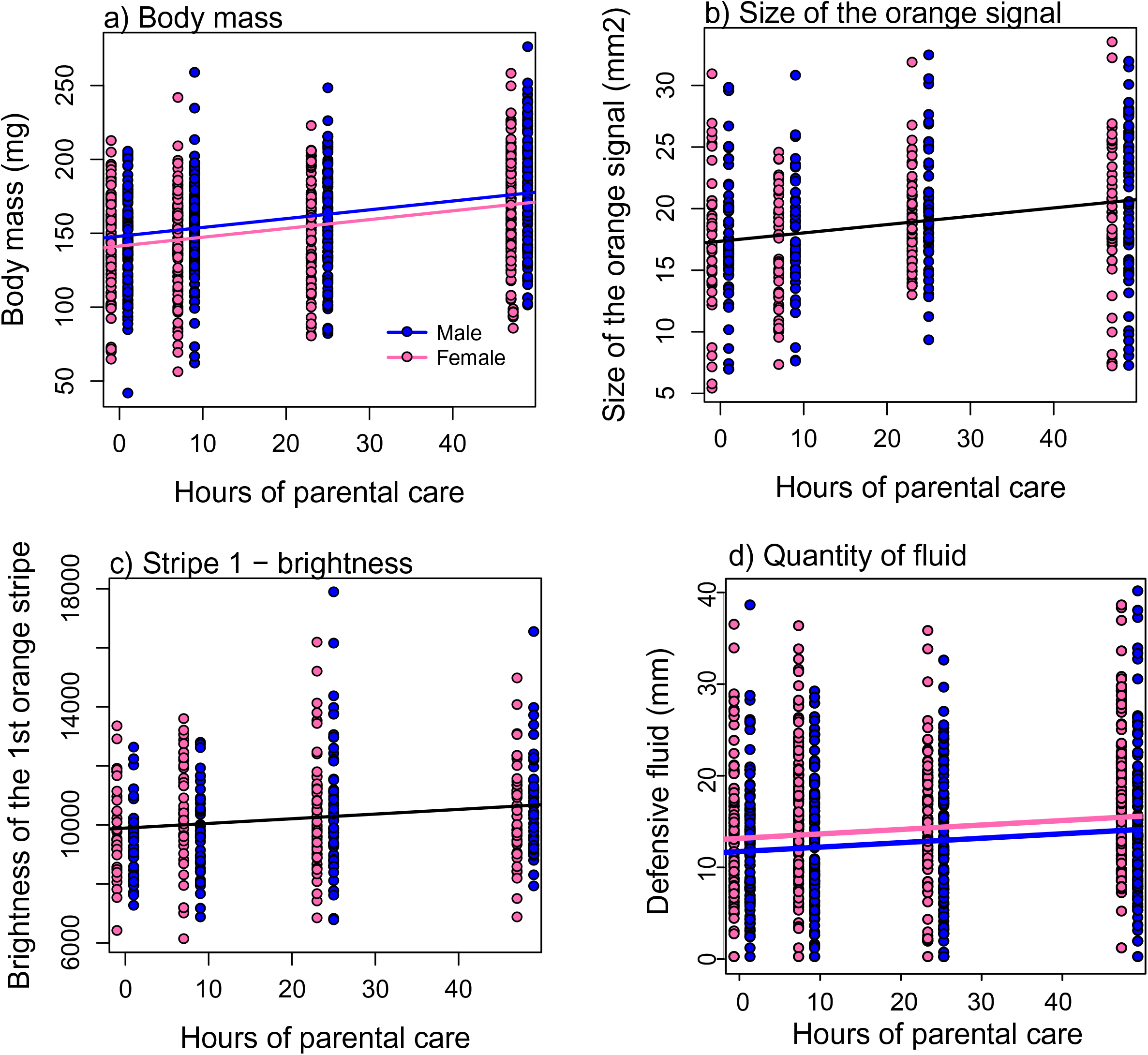
The effect of the duration of care on (a) adult body mass, (b) the total area of the orange signal for males and females, (c) the brightness of stripes and (d) quantity of the defensive fluid produced by males and females. In each case, the fitted lines are predictions from the GLMM.

### Warning signal size

We found that the absolute size of the orange warning signal pattern increased with the level of parental care provided (Type 1 SS: Care duration: F_1,50_ = 8.25, P = 0.006; Fig. 1 b). However, after accounting for elytra size in the model, this effect disappeared (Type 3 SS: Elytra size: F_1,407_ = 430.11, P < 0.001; Care duration: F_1,51_ = 0.32, P = 0.57), suggesting that the effect was driven by the increase in beetle size caused by greater levels of care (Fig. S2). The size of the orange signal was larger in males even after controlling statistically for elytra size (Type 1 SS: Sex: F_1,373_ = 7.68, P = 0.006; Type 3 SS: Sex: F_1,372_ = 3.86, P = 0.050).

Next we investigated whether the relative investment in orange vs. black pigmentation changed with the duration of parental care received as a larva. We found that the proportion of the elytra covered by the orange signal was not affected by the duration of care, but it did increase with the size of the elytra, suggesting that larger beetles invested relatively more into the size of the orange signal than small beetles (Elytra size: F_1,405_ = 20.16, P < 0.001; Care duration: F_1,51_ = 0. 20, P = 0.66, Fig. S2d).

### Warning signal salience

The brightness of the first orange stripe increased with the duration of care received as a larva, independently of the beetle’s size and sex (Care duration: F_1,42_ = 4.91, P = 0.032; Elytra size: F_1,280_ = 2.58, P = 0.11; Sex: F_1,317_ = 0.47, P = 0.49; Fig. 1c). In contrast, the brightness of the smaller, second orange stripe was not affected by the duration of care or the beetle’s sex, but it did increase with the size of the beetle (Care duration: F_1,41_ = 1.47, P = 0.23; Elytra size: F_1,358_ = 4.63, P = 0.032) (Fig. S3 and S4).

The saturation of the first orange stripe was not affected by the duration of care received as a larva, but it did increase with beetle size and was slightly higher in males (Care duration: F_1,40_ = 0.81, P = 0.37; Elytra size: F_1,205_ = 7.24, P = 0.008; Sex: F_1,894_ = 5.29, P = 0.022). The saturation of the second orange stripe also increased with beetle size and, for females, also increased with the duration of care. In contrast, for males, saturation of the second stripe decreased slightly with the duration of care beetles experienced. Males had more saturated second stripes than females (Care duration: F_1,43_ = 0.60, P = 0.44; Elytra size: F_1,345_ = 87.73, P < 0.001; Sex: F_1,460_ = 15.97, P < 0.001; Care duration*Sex: F_1,452_ = 4.90, P = 0.027) (Fig S3 and S4).

The brightness and saturation of the black pattern on the elytra were not explained by sex, elytra size or duration of care or the interactions between any of the variables (F< 3.41, P>0.065).

### Volume and repellence of the defensive fluid

The volume of defensive fluid exuded under threat increased with the duration of care (Type 1 SS: Care duration: F_1,44_ = 6.87, P = 0.012), but, as above, this effect was mainly driven by the effect of parental care on beetle size (Type 3 SS: Care duration: F_1,46_ = 2.85, P = 0.098; Fig. 1d). Females produced more fluid than males and this was independent of size (Elytra size: F_1,324_ = 21.38, P < 0.001; Sex: F_1,380_ = 4.82, P = 0.029; Fig. 1d). The amount of parental care did not significantly affect the repellence of the defensive fluid (Care duration: F_1,213_ = 0.530, P = 0.467), in bioassays involving ants (Fig. S5). However, ants found the defensive fluid produced by males more aversive than that produced by females (Sex: F1,213 = 12.981, P < 0.001). Number of ants drinking the sugar water was positively associated with the number of ants drinking the defensive fluid (F_1,213_ = 197.381, P < 0.001).

### Phenotypic correlation between the warning coloration and chemical defence

We first analyzed the phenotypic correlation between the visual signal (size and color) and the volume of defensive fluid, to test whether the beetle’s coloration was a reliable indicator of its ability to defend itself chemically to potential visual predators (i.e. what predator perceives when attacking the prey). The size of the orange pattern significantly predicted the quantity of the defensive fluid produced (Size of pattern: F_1,276_ = 26.90, P < 0.001; Fig. 2), and this remained the case even if beetle size was retained in the model (Size of pattern: F_1,222_ = 5.97, P = 0.015; Elytra size: F_1,280_ = 3.51, P = 0.061; Sex: F_1,381_ = 5.65, P = 0.018). The effect of duration of care on this correlation was marginal (F_1,46_ = 3.40, P = 0.07) and none of the interaction terms in the model was significant (F<3.49, P>0.06), suggesting that the phenotypic correlation between fluid volume and signal size is independent of the duration of care.

**Figure 2.**
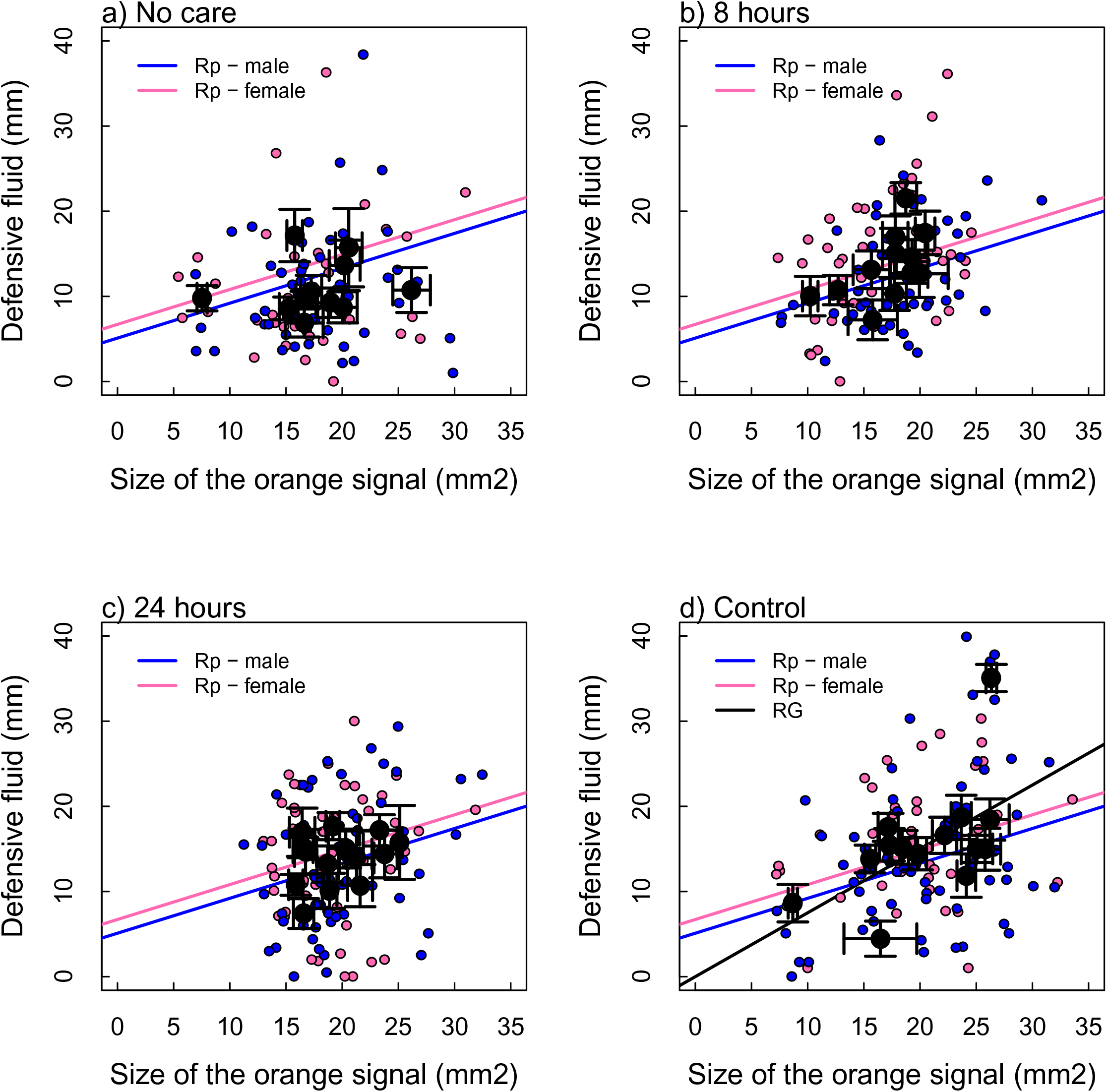
a representation of the phenotypic (Rp) and genetic (RF) correlations between signal size and the quantity of the defensive fluid produced for (a) No care, (b) 8 hours of care, (c) 24 hours of care and (d) control (48 hours of care) beetles. The pink and blue lines represent the phenotypic correlation between the traits for females and males respectively. The phenotypic correlation was not affected by treatment and so the fitted lines from the model with treatment removed are plotted. The family correlation (black line) was estimated using MCMCglmm and was significant for the control treatment only.

Measures of saturation and brightness were not significant predictors of fluid quantity, either as main effects or in interaction with other terms in the models (F<2.84, P>0.09). Therefore, individuals with larger orange patterns produced higher quantities of anal exudate, irrespective of their saturation or brightness. This suggests that the size of the orange pattern could be a reliable signal of an individual’s capacity for chemical defence against predators.

### Family level variation and correlations between the warning coloration and chemical defence

We analyzed the broad sense heritabilities of the component parts of the warning display (Table 1). We found that whereas the broad sense heritabilities of orange pattern size were typically moderate to high in all parental care treatments, the broad sense heritability of the volume of defensive fluid produced was high only in individuals that had full maternal care as larvae.

**Table 1.**
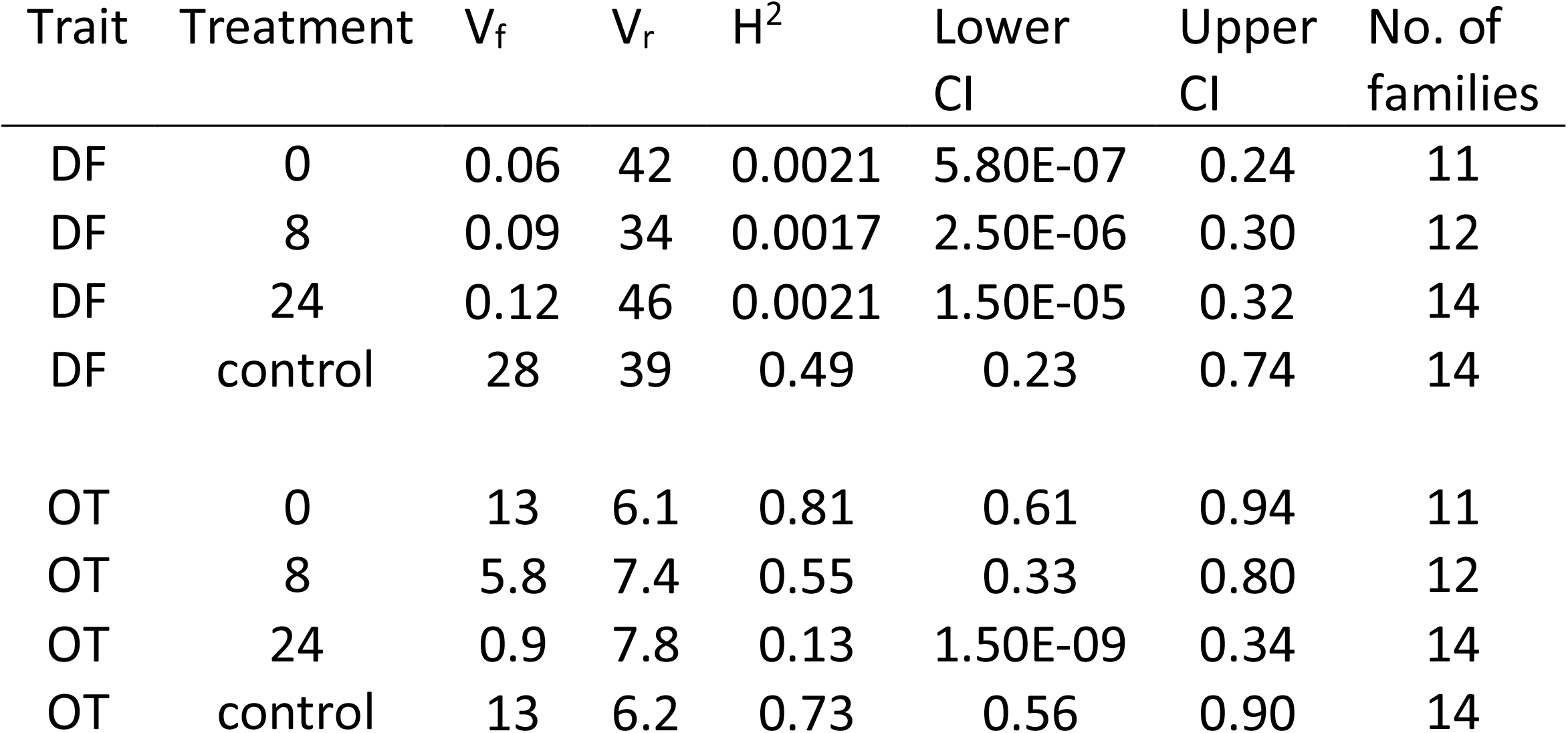
Broad-sense heritabilities (H^2^) for defence fluid volume (DF) and total orange area (OT) for different levels of parental care, calculated from the bivariate MCMC models. V_r_ = residual variance. Genetic variance (additive and non-additive) is estimated using the Family-level variance parameter, denoted V_f_ for clarity. H^2^ is calculated as V_f_/(V_f_+V_r_).

| Trait | Treatment | $V_f$ | $V_r$ | $H^2$ | Lower CI | Upper CI | No. of families |
| --- | --- | --- | --- | --- | --- | --- | --- |
| DF | 0 | 0.06 | 42 | 0.0021 | 5.80E-07 | 0.24 | 11 |
| DF | 8 | 0.09 | 34 | 0.0017 | 2.50E-06 | 0.30 | 12 |
| DF | 24 | 0.12 | 46 | 0.0021 | 1.50E-05 | 0.32 | 14 |
| DF | control | 28 | 39 | 0.49 | 0.23 | 0.74 | 14 |
| OT | 0 | 13 | 6.1 | 0.81 | 0.61 | 0.94 | 11 |
| OT | 8 | 5.8 | 7.4 | 0.55 | 0.33 | 0.80 | 12 |
| OT | 24 | 0.9 | 7.8 | 0.13 | 1.50E-09 | 0.34 | 14 |
| OT | control | 13 | 6.2 | 0.73 | 0.56 | 0.90 | 14 |

We next estimated the family-level (genetic) covariation between the intensity of the orange warning display and the strength of chemical defense, to understand if selection by predators could result in correlated responses in both of these traits. We fitted separate bivariate MCMC models within sets of families receiving different parental care treatments, to estimate the correlation at the family level between defensive fluid and various signal components. Then we compared the family-level correlation coefficient among these models. The strength of the correlations that we measured with these analyses can be attributed to a combination of (1) any genetic correlation between traits, and (2) the common environment shared by members of the same family.

The models returned qualitatively similar estimates for fixed effects (sex and body size) as did the univariate models given above. Family level correlations between defensive fluid volume and brightness were never significantly different from zero (Table 2). Family level correlations between defensive fluid volume and both (1) the total area of orange patterning and (2) the saturation of this orange coloration were also indistinguishable from zero in all families - except broods that received the maximum level of care (Table 2). In these families the correlation with the volume of exudate produced was positive for both measures of the visual display. This means, that under full parental care, the families that on average display larger patches of orange, and which are more saturated in color, also produce higher quantities of defensive fluid.

**Table 2.** Family-level and residual structure in bivariate MCMC models of (a) defensive fluid [DF] and total area of orange [OT], and (b) DF and saturation of the first stripe [SAT], fitted using the MCMCglmm package in R. Values are the posterior mode of each distribution with the upper and lower confidence 95% intervals in parentheses. In each case models were fitted separately for each experimental treatment, and in addition to Family as a random effect, body size and sex were fitted as fixed effects (data not shown).

| Family-level structure |  |  |  |  | Residual structure |  |  |  |
| --- | --- | --- | --- | --- | --- | --- | --- | --- |
| (a) | Family Var (DF) | Family Corr (DF,OT) | Family Cov (DF,OT) | Family Var (OT) | Residual Var (DF) | Residual Corr (DF,OT) | Residual Cov (DF,OT) | Residual Var (OT) |
|  | 0.12 | -0.27 | 0.0047 | 16 | 48 | -0.27 | 0.4 | 6 |
| 0h | (4.9x10<sup>-05</sup>-16) | (-0.88-0.64) | (-10-9.1) | (6.7-55) | (32-60) | (-0.88-0.64) | (-3-4.8) | (4.2-8) |
|  | 0.083 | 0.11 | 0.45 | 7.6 | 35 | 0.11 | 3 | 7.7 |
| 8h | (2.6x10<sup>-05</sup>-16) | (-0.53-0.88) | (-4.5-7.7) | (2.9-26) | (26-49) | (-0.53-0.88) | (-0.79-6.4) | (5.4-10) |
|  | 0.22 | 0.15 | -0.012 | 0.52 | 47 | 0.15 | 2.8 | 8.6 |
| 24h | (0.00036-20) | (-0.79-0.81) | (-3.2-2.4) | (1.6x10<sup>-05</sup>-4.3) | (36-60) | (-0.79-0.81) | (-1.5-6.1) | (6-10) |
|  | 31 | 0.67 | 10 | 16 | 41 | 0.67 | 0.86 | 6.7 |
| Control | (10-100) | (0.00058-0.91) | (-3.9-37) | (5.7-38) | (31-55) | (0.00058-0.91) | (-3.1-4) | (5-8.8) |
| (b) | Family Var (DF) | Family Corr (DF,SAT) | Family Cov (DF,SAT) | Family Var (SAT) | Residual Var (DF) | Residual Corr (DF,SAT) | Residual Cov (DF,SAT) | Residual Var (SAT) |
|  | 0.15 | 0.38 | -6.5x10<sup>-05</sup> | -9.5x10<sup>-07</sup> | 40 | 0.38 | 0.0062 | 0.00029 |
| 0h | (3.6x10<sup>-06</sup>-17) | (-0.77-0.89) | (-0.0075-0.012) | (4.2x10<sup>-12</sup>-6.9x10<sup>-05</sup>) | (33-61) | (-0.77-0.89) | (-0.03-0.025) | (0.00021-0.00039) |
|  | 0.09 | -0.081 | 6.2x10<sup>-06</sup> | 5.3x10<sup>-06</sup> | 39 | -0.081 | 0.0086 | 0.00056 |
| 8h | (0.00014-17) | (-0.95-0.77) | (-0.012-0.0096) | (1.3x10<sup>-11</sup>-8x10<sup>-05</sup>) | (26-49) | (-0.95-0.77) | (-0.028-0.034) | (0.00037-7x10<sup>-04</sup>) |
|  | 2.4 | -0.75 | 4.1x10<sup>-05</sup> | 1.5x10<sup>-06</sup> | 47 | -0.75 | 0.0066 | 0.0011 |
| 24h | (0.00021-19) | (-0.99-0.54) | (-0.035-0.014) | (1.3x10<sup>-09</sup>-0.00034) | (33-60) | (-0.99-0.54) | (-0.042-0.053) | (0.00082-0.0014) |
|  |  | 0.92 | 0.032 | 8.2x10<sup>-05</sup> | 41 | 0.92 | -0.012 | 0.00039 |
| Control | 33 (13-90) | (0.048-1) | (-0.011-0.15) | (1.1x10<sup>-07</sup>-0.00041) | (29-53) | (0.048-1) | (-0.035-0.019) | (0.00027-0.00051) |

## DISCUSSION

Our results show that parental care can have long-term effects on the development and evolution of aposematic displays. First, we show that the greater the level of care received by larvae, the more salient is the visual warning signal to predators in adulthood (brightness in both sexes and color saturation in females). Furthermore, individuals that received more care matured into larger adults, displayed correspondingly larger warning signal patterns and produced greater volumes of defensive fluid when subjected to simulated attack. Previous studies have shown that parental care defends juveniles against predators, by offering the protection of a nest (37); or through guarding behaviour (38); or by chemically defending offspring in diverse ways (39–43). A key new insight from our study is that parental effects can offer protection against predators that endure even when the offspring become adults.

Second, we found a positive phenotypic correlation between the size of an individual’s visual warning display and the extent of its chemical defense, which was marginally affected by the level of care received during development. From a potential predator’s perspective individual beetles with large orange patterns are likely to exude larger volumes of defensive fluid than individuals with smaller orange patterns. This means that predators should avoid beetles with large warning signals as they are likely to have a more effective chemical defense, thus potentially placing a positive selection pressure on the size of the signal (44, 45).

However, the reliable information content of the warning display can only be maintained under predator-induced selection if its components are genetically correlated i.e. selection on either of the components (e.g. pattern size) results in a correlated response also in the other component (e.g. chemical defence). To understand how parental care might modulate the strength of this correlation, we determined the correlation between the visual components of the display and the chemical defenses after statistically eliminating other environmental sources of variation. With these analyses, we found that there was a genetic correlation between the visual components of the display and the strength of the chemical defense only in individuals that received full maternal care as larvae (3, 6). This suggests that, under conditions where there is a reduced supply of care, selection on the size of the warning signal would not result in a correlated response in the quantity of the chemical defense over evolutionary-time. Thus, the extent of parental care supplied within the prey species can modulate the response to selection by predators on the information content of its warning display.

We can think of two explanations for a decay in the correlation between the visual display and the strength of the chemical defenses at low levels of care. First, low levels of maternal care are more stressful and increase environmental variation in chemical defenses, thus reducing broad sense heritability (46, 47). Accordingly, our results suggest that within reduced care families (poor condition), we had some individuals who may have monopolised the resource (or who were just better at self-feeding), and therefore end up with enough resources to become large and have a large orange pattern but still be able to invest in high volumes of defensive fluid. Some of their siblings may still have had relatively large orange signals (most likely due to its high heritability) but due to stressful conditions (increased competition for food or poor ability for self-feeding) could not invest in high volumes of defensive fluid (Fig. 2). This higher within-brood variation could also explain why we found a positive phenotypic correlation between the signal and defence irrespective of parental care treatment, but on the family level only under full care treatment.

Secondly, perhaps our measures of broad sense heritability for chemical defenses are driven solely by parental effects. For example, parents can use selective feeding and provide extra nutrients to smaller larvae, thus reducing within-brood variation under full care in comparison to those receiving reduced care or no care at all. Because parents vary in quality, the absolute amount of resources each larva within a brood receives is likely to vary also with the quality of the parent resulting in family-level variation also under full care in aposematic traits. This explanation is consistent with previous work showing that the early-life environment experienced by tiger moths influences the potency of their chemical defenses (33).

Unexpectedly, we found sexual dimorphism both in the salience of the visual display (males were more saturated in orange coloring than females) and in the chemical defenses (females produced more fluid in response to attack, while male exudates were more noxious to ants). These sex differences might simply reflect different sex-specific roles in reproduction and parental care (18). Previous work has found that females allocate more effort to the maintenance of the carcass during biparental care, which might involve producing higher quantities of anal exudate to spread on the carcass. By contrast, males put more effort defending the carcass against other beetles and other competitors such as ants and flies (48–51). This might explain why their exudates are more repellent.

More generally, our results provide new insights into the evolution of warning displays. Classically, research on warning signals has focused on determining the evolutionary outcomes of predator-prey interactions, or the selective constraints imposed by the ecological or physiological condition of both the prey and the predator (52). Recently it has become evident that the predator’s social environment additionally influences warning signal evolution, by changing patterns of selection on prey. For example, social information received from other conspecifics affects how predators choose their prey and what kind of prey they should avoid (53). Our study suggests that the social environment experienced by prey is also important, because it determines the response to selection on the honesty of the warning display imposed by predators. Many aposematic insect species such as wasps, bees, Lepidopteran larvae, and sawflies (54), as well as aposematic vertebrates (16), inhabit a social environment of conspecifics as a result of their gregarious life-style, or parental care, or eusociality. Our work suggests that the prey’s social environment could be generally important in determining the pace and direction of evolutionary change in warning displays, in response to selection from predators.

## MATERIALS AND METHODS

### N. vespilloides colony

We used the same outbred laboratory population of burying beetles established in 2005 at Cambridge University as described in (18). Details of animal husbandry are included in the Supplementary Information.

### Experimental procedure

All treatments were run between July 2010 and September 2011. We set up one hundred boxes, each with a carcass and a pair of adults. All males were removed from the boxes ~53 hours after pairing, i.e. after egg laying but before hatching had commenced. Broods of the widowed females were haphazardly divided among the following four treatments: ‘0 h’ maternal care: mothers removed at hatching; ‘8 h’ maternal care: mothers removed from the boxes 8h after the larvae hatched; ‘24 h’ maternal care: mothers removed from the boxes 24 h after the larvae hatched; ‘> 48 h’ maternal care: mothers retained until larvae dispersed from the carcass to pupate. Larvae typically hatched 71.28 h +-1.47 SE after pairing (55) and therefore boxes were checked every ca. 5 h from 55-96 h after pairing to determine the time of hatching. When larvae dispersed, eight days after pairing their parents, they were transferred to separate boxes, to pupate. In total, 97 pairs bred successfully, yielding 24-25 families per treatment.

At eclosion, when individuals had developed the typical black and orange coloration their chemical defences were measured as described in (18) and in the Supplementary Information. To quantify the chemical defence, we held each beetle and gently tapped the abdomen from the ventral side. We collected the resulting droplet of fluid in a capillary tube, and measured the quantity produced. We aimed to collect samples from five females and five males per family, when the number of offspring per family was sufficiently large. Altogether we sampled 2-10 individuals per family (mean = 8.37, SD = 2.04) yielding 812 samples in total. After collecting these samples, individuals were sexed, weighed and stored in a freezer at −20 ° C for color analysis.

To estimate variation in the toxicity of the defensive secretion among parental care treatments, we conducted standard bioassays with ants (*Formica spp*) similar to those described previously (18) and in Supplementary information. We tested the effectiveness of the anal exudate in deterring wood ants (*Formica* spp.) by offering them either a 10% exudate solution (10 % anal exudate / 90 % sugar water) or a 10 % control solution (10% water / 90% sugar water). We know from our previous work that this concentration of exudates is sufficiently potent to change ant behaviour (18). The samples we collected to estimate the volume of the fluid produced by individual beetles were pooled together into Eppendorf tubes, by maternal care treatment, and by sex (i.e. we had one tube for each sex, per maternal care treatment). Each ant nest (N=9) was tested three times (N=3) with pooled fluid samples collected from each sex (N=2) within each maternal care treatment (N=4). (N = 9 x 3 x 2x 4 = 216 possible trials of which 213 yielded data).

To measure the pattern size and color, we photographed beetles using a calibrated Fuji IS digital camera, which records both ultraviolet and human visible signals. From the photographs we measured the size of the orange markings on the elytra as well as brightness and saturation of their orange and black colour to potential bird predators. For pattern size measurements we used Image J –program and color measurements were done by using the Image Calibration and Analysis Toolbox (56), using the method described in (18).

### Statistics

All phenotypic traits i.e. body size, the size and color (saturation and brightness) of the orange signal and the quantity and quality of the defensive fluid, were analysed using linear mixed effects models using the *lme4* and *lmerTest* packages (57, 58) in the *R* statistical package version 3.2.2 (R Core Team, 2013). For every model, the *family* from which the beetles were derived, nested within *treatment* was included as a random effect. For the fixed effects, initial data exploration suggested that most traits of interest increased linearly with the duration of care. Therefore, treatment was coded as a continuous variable corresponding to the *Duration of care* in hours.

We considered the effects of *Duration of care* and *Sex* on the variation in body mass and elytra size. For the color traits and defense fluid quantity, we additionally considered the effects of *Elytra size* although the results were very similar if we included *Body mass* in its place. In all cases we fitted the maximal model with all interaction terms included and used stepwise deletion until we were left with significant terms only. The residuals from each model were visually examined for deviations from normality. The saturation of the orange stripes showed significant deviations from normality and so were subjected transformation using the transformTukey function in the *Rcompanion* package (59).

We also looked at the phenotypic relationships between the signal (size and color) and the quantity of defensive fluid to see if the signal was a reliable indicator of defensive capability on individual level. For these models, the quantity of defensive fluid was the response variable and in addition to the *Duration of care*, *Sex* and *Elytra size*, we considered the effects of *Signal size* and color (*Saturation* and *Brightness*) in separate models.

To test the effect of *Duration of care in hours* and *Sex* of the beetle on the deterrence of defence fluid against ants (number of ants drinking the exudate), we used a general linear mixed model. Treatment was again coded as a continuous variable. Number of ants drinking the sugar water was included as a covariate and *Ant nest* nested with *Trial number* were included as a random factor. Non-significant parameters were omitted from the final model (general protocol; p-value > 0.05, smallest omitted significance p = 0.565).

To test the phenotypic relationship between the warning display and quantity of chemical defense (i.e. what predator perceives when attacking the prey), we used a General Linear Mixed model in R. As a response variable we had quantity of defense fluid and elytra size, size of the orange pattern, sex and parental care treatment as a fixed effect. Family was included as a random effect under the treatment.

### Quantitative genetic analyses

To estimate the family correlation (which includes the genetic correlation) and the broad-sense heritabilities of each trait, we used the MCMCglmm package in R to fit bivariate linear mixed models in a MCMC framework. In each case the bivariate response variable was a combination of “defense fluid volume” and one of (1) size of the orange pattern, (2) saturation of the orange pattern, and (3) brightness of the orange pattern (double cone sensitivity). For each bivariate combination of responses, we fitted four separate models, in each case using subsets of families receiving different levels of parental care. In each model we fitted sex and body size (mg) as fixed covariates, plus a family-level random effect to estimate the family correlation.

Family level correlations include additive genetic, dominance, permanent environment, maternal, and paternal effects. We calculated the family correlation as:

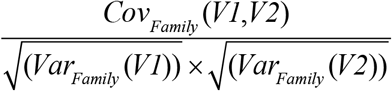

Broad-sense heritability in the response traits for each model was calculated as Var_Family_ / Var_Total_

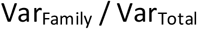

Where Var_Total_ = Var_Family_ + Var_Residual_.

We fitted each model using parameter-expanded priors for the residual variance structure with nu = 2 (T. Houslay, pers. comm.). We also ran the models under a range of alternative priors for the residual component of the model (e.g. informative priors assuming equally apportioned variance, inverse-Wishart distributed priors) to check for robustness to starting assumptions. Three independent chains were run for each model and these were all checked by eye for autocorrelation and convergence with no issues.

## ACKNOWLEDGEMENTS

This study was funded by the Academy of Finland via the projects no 136387, 257581 (C.L.). G.B. was funded by Marie Curie Intra-European Fellowship 252120 LIFHISBURBEE. S.C.C. was supported by a Natural Environment Research Council grant (NE/H014225/2). R.M.K. was supported in part by the European Research Council (ERC) Consolidators grant 310785 BALDWINIAN_BEETLES and by a Wolfson Merit Award from the Royal Society. We are grateful to Amy Backhouse for helping with the maintenance of lab population; to Martin Stevens and Jolyon Troscianco for helping with the color analyses; and to Tom Houslay for helping with the statistical analyses.

