## Supplementary Information for "Parental care shapes evolution of aposematism and provides lifelong protection against predators"

SUPPLEMENTARY FIGURES

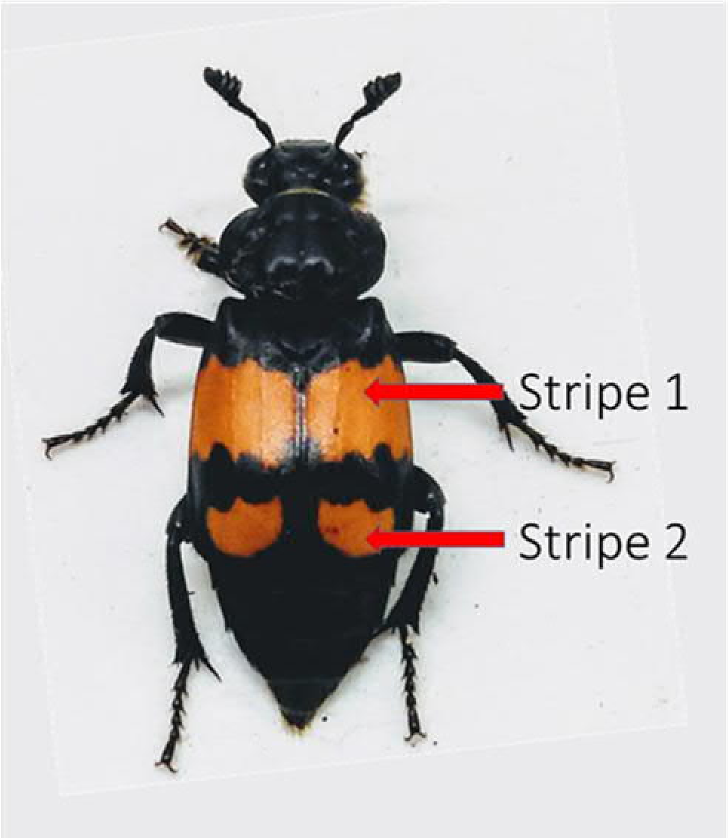


Figure S1 – A Nicrophorus vespilloides adult showing two stripes of conspicuous orange markings on its elytra.

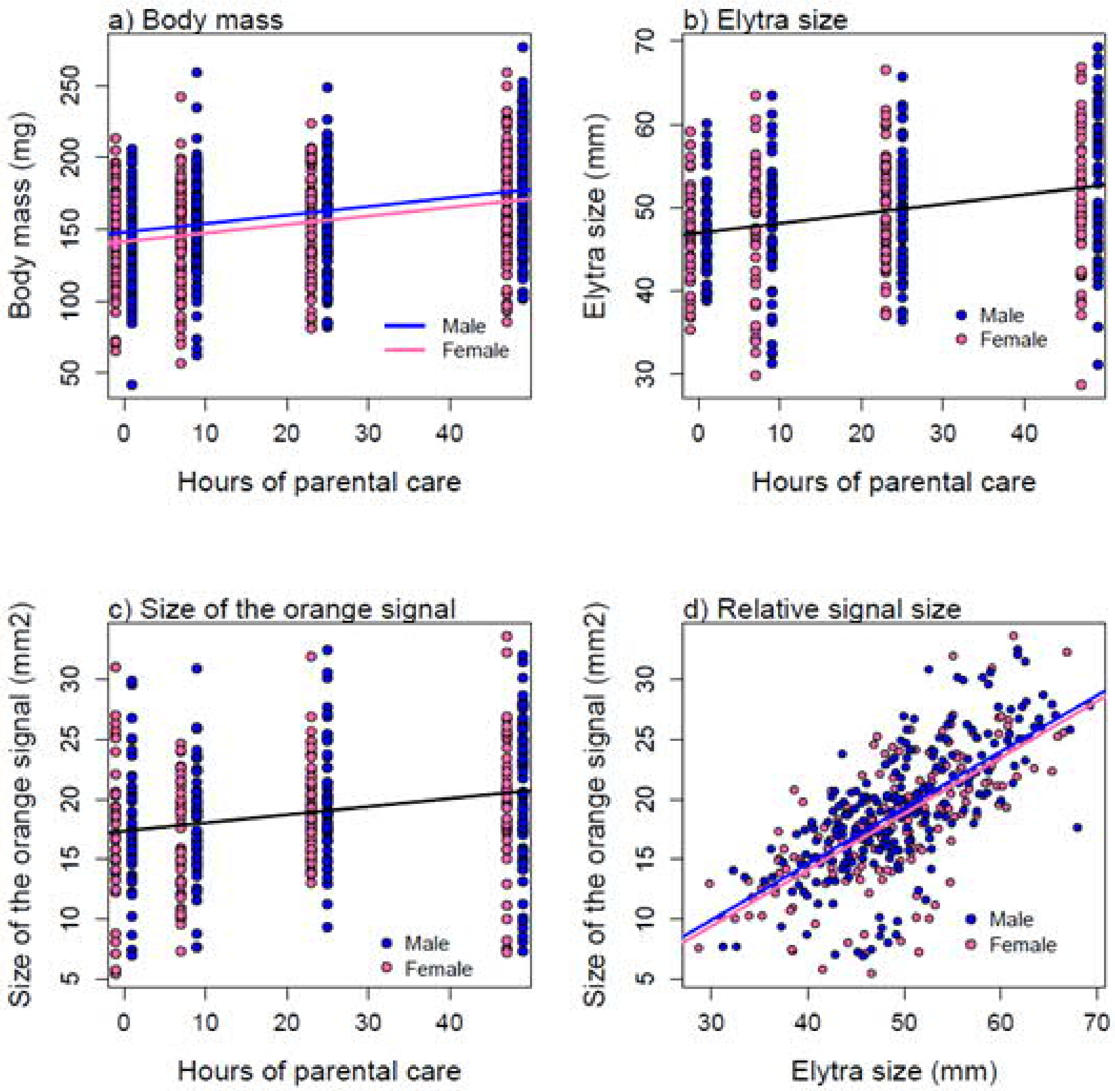


Figure S2 - The effect of the duration of care on (a) adult body mass, (b) the size of the elytra and (c) the total area of the orange signal for males and females. The relationship between signal size and the size of the elytra is also shown (d). In each case, the fitted lines are predictions from the GLMM.

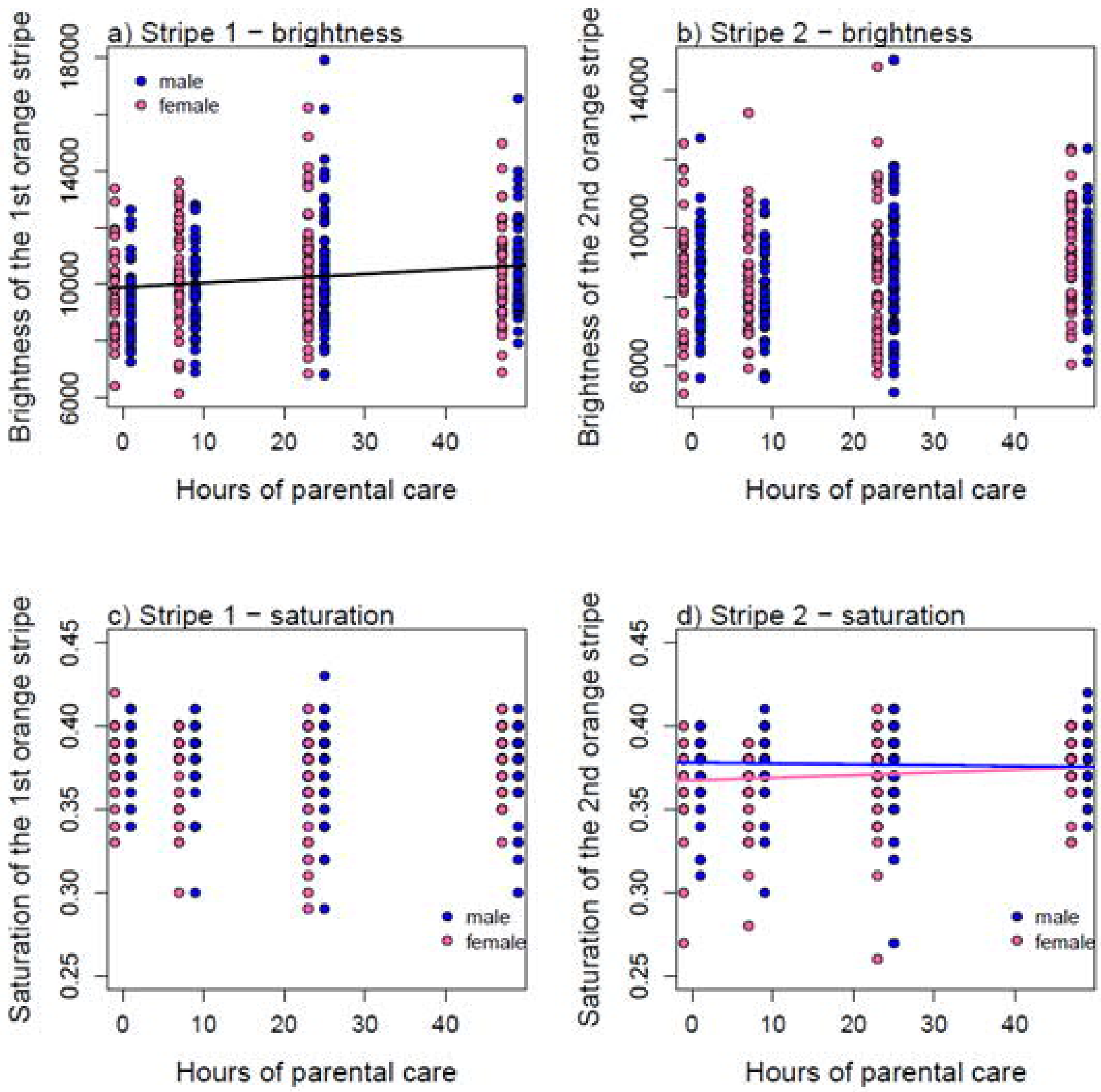


Figure S3 - The effect of the duration of care the brightness of stripes 1 (a) and 2 (b) and the saturation of stripes 1 (c) and 2 (d) for males and females. In each case, the fitted lines are predictions from the GLMM.

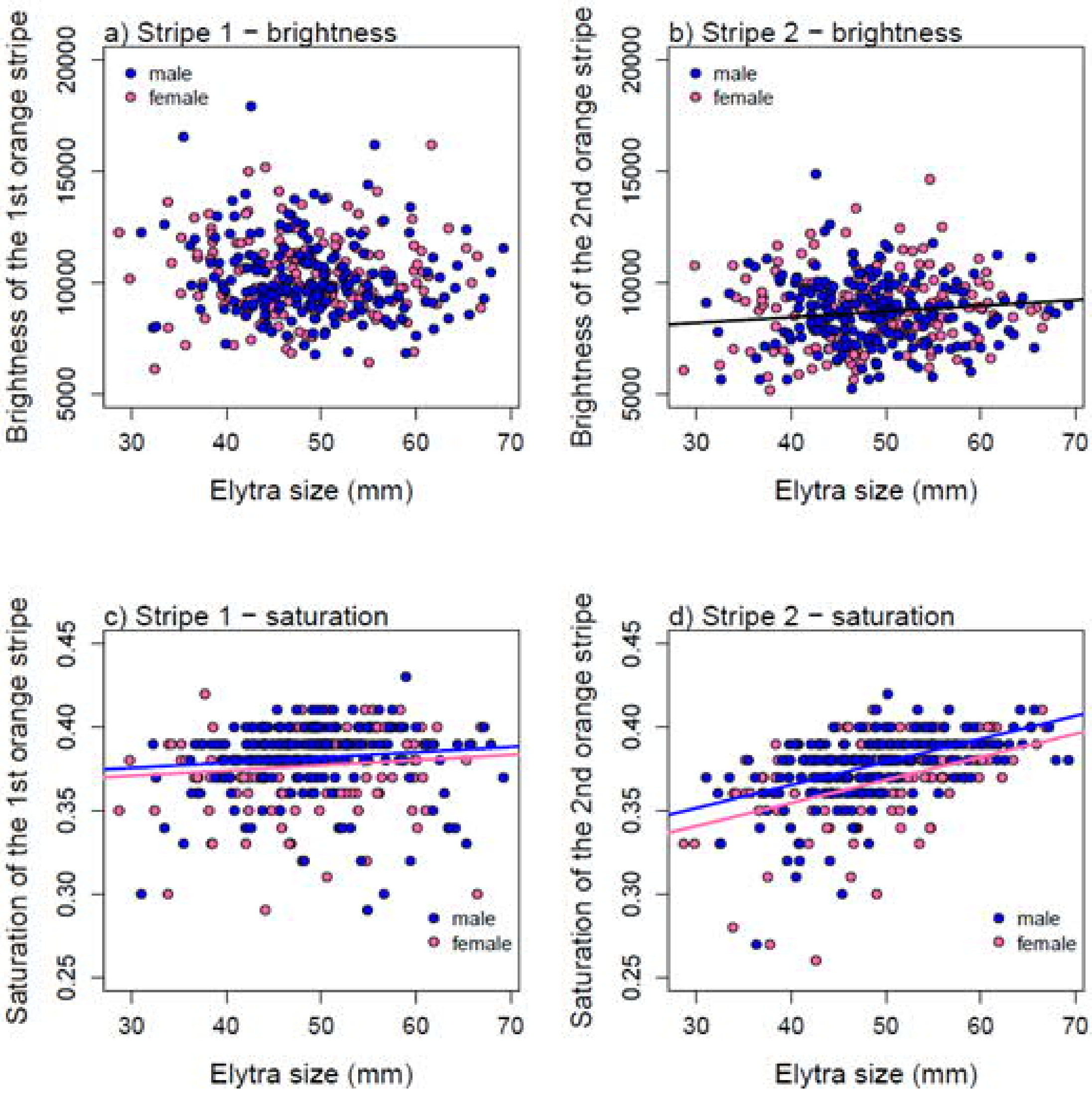


Figure S4 - The relationship between the size of the elytra and the brightness of stripes 1 (a) and 2 (b) and the saturation of stripes 1 (c) and 2 (d) for males and females. In each case, the fitted lines are predictions from the GLMM.

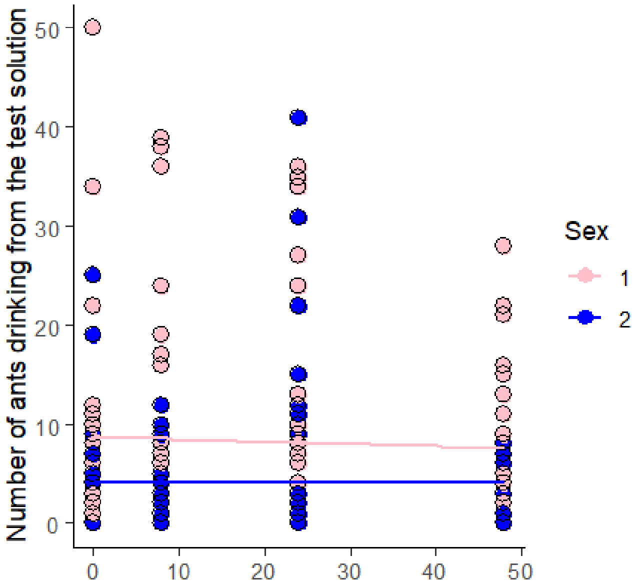


Figure S5 - The effect of the duration of care on repellence of the defensive fluid produced by males (blue) and females (pink).

### SUPPLEMENTARY METHODS

#### N. vespilloides colony

We used the same outbred laboratory population of burying beetles established in 2005 at Cambridge University as described in (1). Individual adults were housed in moist soil in plastic boxes (12x8x2 cm) and fed minced beef twice weekly. Boxes were kept at a constant temperature of 21 °C on a 16h:8h light:dark cycle. For breeding, unrelated beetles were paired in plastic boxes (17x12x6 cm) half-filled with moist soil. Each pair was provided with a freshly thawed mouse carcass (21.94 ± 0.33 SE g, range 15-35g) and the breeding box placed in a dark cupboard to simulate underground conditions. Larvae dispersed from the carcass ca. 8 days after pairing and eclosed as adults ~3 weeks later. Sexual maturity was reached 1-2 weeks after eclosion.

Effect of maternal care on variation in quantity and toxicity of chemical defences

At eclosion, when individuals had developed the typical black and orange coloration, we simulated an attack by a predator and measured how much defence fluid was produced from the anus by both males and females from each of the different parental care treatments. Specifically, we held each beetle and gently tapped the abdomen from the ventral side. We collected the resulting droplet of fluid in a capillary tube, and measured the quantity produced. We aimed to collect samples from five females and five males per family, when the number of offspring per family was sufficiently large. Altogether we sampled 2-10 individuals per family (mean = 8.37, SD = 2.04) yielding 812 samples in total. After collecting these samples, individuals were sexed, weighed and stored in a freezer at -20 ° C for colour analysis (see below).

To estimate variation in the toxicity of the defensive secretion among parental care treatments, we conducted standard bioassays with ants (Formica spp) similar to those described previously (1, 2). The burying beetle’s aposematic display is likely to have been selected by visual predators, such as scavenging birds (see (1), and references therein). Nevertheless, ants and burying beetles are also rivals for carrion and the burying beetle’s chemical defences are likely to deployed against ants (3). The defensive fluid of burying beetles contains several defensive compounds that all are likely to contribute their unprofitability synergistically (1, 4). This means the extent to which ants find the chemical defences of the burying beetles aversive provides a good biomarker of this fluid’s toxic potency.

The samples we collected to estimate the volume of the fluid produced by individual beetles were pooled together into Eppendorf tubes, by maternal care treatment, and by sex (i.e. we had one tube for each sex, per maternal care treatment). These pooled samples were stored in a freezer (at -20 ° C), until they were analysed. For testing, the samples were thawed and then diluted with a 20 % sugar solution (20% sugar, 80% water) to make the solution potentially attractive to ants. We tested the effectiveness of the anal exudate in deterring ants by offering them either a 10% exudate solution (10 % anal exudate / 90 % sugar water) or a 10 % control solution (10% water / 90% sugar water). We know from our previous work that this concentration of exudates is sufficiently potent to change ant behaviour (1).

We performed tests with the 10% exudate and control solutions on 9 different ant (Formica spp.) nests in central Finland (62 °N, 26° E) in sunny and warm weather (15-20°C). All tests were run within a 1 week period in August 2011. To standardize the potential variation in activity and ant traffic among ant nests, we presented ants simultaneously with droplets of exudate and control solutions. In the vicinity of each nest we chose a spot on the trail where ant traffic was about 10 to 20 individuals/minute. We put 10 μl of both solutions close to each other (<2 cm) on a transparent, sterilized plastic circle (4 cm in diameter) and offered it to the ants. We repeated the assay three times per nest on three different ant trails, changing the sequence in which the two treatments were presented between repetitions. After presentation, we counted the number of ants drinking from the different solutions every minute for the next 10 minutes (1, 2), and started this observation period when the first ant worker encountered either of the two droplets. Each ant nest (N=9) was tested three times (N=3) with pooled fluid samples collected from each sex (N=2) within each maternal care treatment (N=4). (N = 9 x 3 x 2x 4 = 216 possible trials of which 213 yielded data).

Effect of maternal care on variation orange elytral patterning and colour

Nicrophorus vespilloides adults have two stripes of conspicuous orange markings on their elytra (Figure 1), which we have previously shown to act as an aposematic signal (1). Frozen individuals were photographed using a calibrated Fuji IS digital camera, which records both ultraviolet and human visible signals. From the photographs, the size of the elytra and of the orange stripes were measured with Image J similar to (1). In addition, we chose approximately similar sized square shaped samples (region of interests, ROIs) from the left side of the beetle's elytra from both of the orange stripes as well as from the black markings. The hue and brightness of the orange and black markings were analysed with the Image Calibration and Analysis Toolbox (5), using the method described in (1). We calculated saturation values (color richness), and brightness (double cone sensitivity) after (6) for the ROIs of the first and second orange stripes and black pattern.

Signal size measurements were taken from 416 individuals across 51 families, with 11-14 families per treatment (Figure 1). Altogether we sampled 2-10 individuals per family (mean = 8.16, SD = 2.16). Colour values of the signal were collected from 378 individuals across 43 families, with 9-12 families per treatment. Altogether we sampled 3-10 individuals per family (mean = 8.79, SD = 1.60).
